# The [FeFe]-hydrogenase *So*HydAB of *Shewanella oneidensis* – An Atypical M2(D)-type Enzyme that Drives Formate Oxidation

**DOI:** 10.64898/2026.10.15.759383

**Authors:** Florian Riemer, Max A. Klamke, Anna-Maria Wandinger, Thomas Happe, Sven T. Stripp, Eckhard Hofmann

**Affiliations:** Structural Biology and Biospectroscopy, Faculty of Biology and Biotechnology, Ruhr University Bochum, Universitätsstraße 150, 44801 Bochum, Germany; Spectroscopy & Biocatalysis, Institute of Chemistry, University of Potsdam, Karl-Liebknecht-Straße 24-25, 14476 Potsdam, Germany; Photobiotechnology, Faculty of Biology and Biotechnology, Ruhr University Bochum, Universitätsstraße 150, 44801 Bochum, Germany

## Abstract

The gamma-proteobacterium *Shewanella oneidensis* harbours one [NiFe]- and one [FeFe]-hydrogenase. This paper presents the first biochemical characterisation of *So*HydAB, a M2(D)-type [FeFe]-hydrogenase consisting of a small subunit featuring a twin-arginine-targeting motif (TAT) and a large subunit harbouring the active site cofactor and two accessory iron-sulfur clusters. There are three main reasons to investigate this enzyme. Firstly, *S. oneidensis* is a highly studied, metabolically complex, and widely utilised organism, especially in the context of wastewater treatment, CO_2_ conversion, and microbial fuel cells, including H_2_-producing systems. Advanced understanding of its hydrogen metabolism can improve biotechnological applications. Secondly, *S. oneidensis* has been a mainstay of hydrogenase research for decades as its enzymes HydE, F and G are the most widely studied and utilised maturases – enzymes assembling the active site cofactor of [FeFe]-hydrogenases. Lastly, while enzymes like *Nd*HydAB from *Nitratidesulfovibrio vulgaris* (formerly *Dd*HydAB, *Desulfovibrio vulgaris*) with high similarity to *So*HydAB have been characterised extensively in the past, the metabolic context of the enzyme in *Shewanella* is different from *Nitratidesulfovibrio*. While *Nd*HydAB receives electrons from ferredoxin, *So*HydAB accepts electrons from the menaquinone pool of the outer membrane while being a soluble enzyme in the periplasm. This is facilitated by interaction with a membrane-integrated cytochrome b transcribed from the same operon. Oxidation of the menaquinone pool drives oxidation of formate toward CO_2_ via formate dehydrogenase. This study therefore lays the groundwork for understanding H_2_-dependent CO_2_ turnover, a new type of metabolic niche for [FeFe]-hydrogenases, while simultaneously improving our understanding of a widely used platform organism.

## Introduction

*Shewanella oneidensis* is a gamma-proteobacterium first isolated from a sediment from lake Oneida^1^. The type strain MR-1 was originally named for its ability to reduce manganese, although today the acronym is often expanded to “metal-reducing”, as *S. oneidensis* is able to reduce a broad array of metals, including chromium^2^, palladium^3^, uranium^4^ and many others^5^. This is one example of *Shewanella*’s metabolic flexibility. *Shewanella oneidensis* is a facultative anaerobe, can produce nanowires to transfer electrons directly to oxidised metal surfaces^6^, secrete flavin-based electron mediators^7^, and has a wide range of flexible and redundant electron transport chains^8^. Beside this, MR-1 also produces hydrogen gas (H_2_) in the absence of exogenous electron acceptors^9^. All of these properties, taken together with the non-pathogenicity of MR-1 have made this bacterium a popular choice for biotechnological applications such as wastewater treatment, microbial fuel cells, or biohydrogen production^10^. The hydrogen metabolism of *S. oneidensis* has been well characterised physiologically^9,11^ but a comprehensive biochemical characterisation of the responsible enzymes has not been published^12,13^.

In bacteria, archaea, and algae the enzymes responsible for hydrogen turnover are referred to as hydrogenases^14^. Among these, [NiFe]-hydrogenases are mostly active in H_2_ oxidation while [FeFe]-hydrogenases typically catalyse both H_2_ oxidation and H_2_ evolution^15^. The central enzymes in the hydrogen metabolism of MR-1 are one [NiFe]- and one [FeFe]-hydrogenase^9^. The [NiFe]-hydrogenase HyaAB retains activity in the presence of trace amounts of O_2_, contributes the larger share to total H_2_ evolution and was experimentally confirmed to be active in H_2_ oxidation *in vivo*^9^. The [FeFe]-hydrogenase contributes a smaller share of H_2_ evolution, is only expressed once O_2_ is completely undetectable in a culture, and has not been shown to be able to oxidise H_2_^9^. Despite this seemingly unflattering comparison, *Shewanella* is well known in [FeFe]-hydrogenase research. The assembly of the cofactor of [FeFe]-hydrogenase, the so-called H-cluster, requires specialised enzymes referred to as “maturases”^16^. HydE, HydF, and HydG from *S. oneidensis* are model enzymes for [FeFe]-hydrogenase maturation^17,18^. They have also been widely used to produce functional hydrogenase in heterologous expression systems lacking endogenous maturases^19^.

The [FeFe]-hydrogenase of MR-1 consists of one large subunit HydA, featuring the active site and two accessory [4Fe-4S] clusters (F-clusters) as well as a small subunit HydB with a TAT motif for transport into the periplasm. This makes *So*HydAB a member of the M2(D)-type of hydrogenases^20,21^. The [FeFe]-hydrogenase of *Nitratidesulfovibrio vulgaris* (*Nd*HydAB, previously abbreviated DdH or *Dd*HydAB) is the only other hydrogenase of this type that has been extensively studied on a molecular level^22–25^.

The genes *hydA*, *hydB*, *hydE*, *hydF*, and *hydG* are organised in a single operon in MR-1^26^. This operon also contains genes coding for a small protein of unknown function designated *hydX* and a cytochrome b protein designated *hydC* (Fig.1A). HydC has been shown to be an essential part of hydrogen turnover in various bacteria^27^ including MR-1 where it is presumed to deliver electrons from the menaquinol pool to HydAB^11^. Molecular details regarding the interaction of [NiFe]- or [FeFe]-hydrogenase with HydC are not known. It should be noted that in the older literature *hydC* is often referred to as *fdh* as the γ-subunit of formate dehydrogenase (see below) is also a cytochrome b protein.

**Figure 1:**
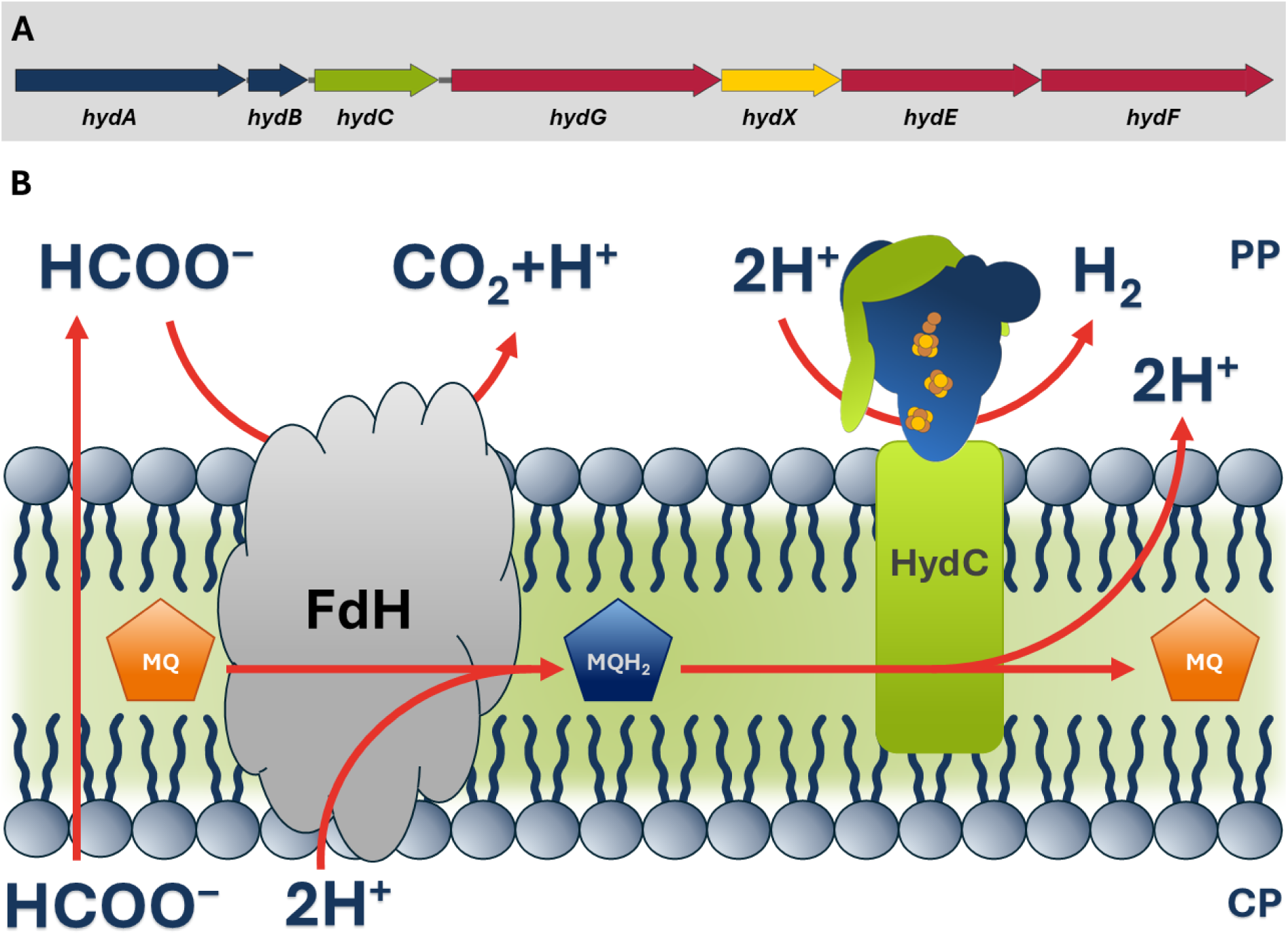
**(A) *hyd*-Operon of *Shewanella oneidensis* MR-1**. The genes *hydA* and *hydB* (in blue) encode the subunits of the hydrogenase. *hydC* (green) codes for a cytochrome b responsible for supplying the hydrogenase with electrons. The maturase machinery (red) consists of *hydG*, *hydE* and *hydF*. The operon also contains a gene for a small protein of unknown function termed *hydX*, shown in yellow. **(B) Simplified representation of the metabolic context of *So*HydAB.** Formate (HCOO^−^) is exported by FocA (protein not shown). Formate dehydrogenase (FdH) oxidises formate to CO_2_. Inside the membrane, two electrons are transferred to menaquinone (MQ) to form menaquinol (MQH_2_) and relayed to HydAB by electron transfer subunit HydC. The protons for the reduction of MQ are taken from the cytoplasm (CP) and released into the periplasm (PP) upon oxidation of MQH_2_. NB: *S. oneidensis* has three FdHs, all of which are periplasmic, membrane-associated multienzyme complexes.

The metabolic reason for H_2_ production in *S. oneidensis* is believed to be the maintenance of proton motive force (PMF) and ATP synthesis^28^. When MR-1 grows anaerobically and in the absence of exogenous electron acceptors like fumarate, it converts pyruvate into formate via pyruvate-formate-lyase^29^. Formate is an important single-carbon (C1) electron carrier^30^; however, the accumulation of formate leads to a drop of the cytoplasmic pH value. To counteract this acidification, MR-1 exports formate to the periplasm via pH-dependent transporter FocA^31^, where it is oxidised toward CO_2_ by formate dehydrogenase (FDH), the transmembrane enzyme that reduces menaquinone (MQ) to menaquinol (MQH_2_) in turn^28,29^. CO_2_ leaves the cell or is recaptured by the help of carbonic anhydrase^32^; however, formate oxidation releases protons in the periplasm that the bacterium needs to dispose of to balance acidification.

This is achieved by soluble [FeFe]-hydrogenase *So*HydAB, located in the periplasm and catalysing the reduction of protons toward H_2_, which leaves the cell as off-gas (Fig. 1B). Protons, in this context, act as “exogenous” electron acceptor. *So*HydAB receives electrons from membrane-associated HydC that re-oxidizes the MQ pool, previously reduced by FDH^11^. Note that during MQ/MQH_2_ conversion, a pair of protons is exported from the cytoplasm^33^. This seemingly zero-sum metabolic shunt disposes of electrons, maintains PMF and ATP synthesis, and constitutes an adaptation to life on the metabolic edge in the nutrient-poor sediments that MR-1 typically inhabits^34^. While this situation constitutes a novel metabolic niche for [FeFe]-hydrogenases, similar pathways have been described involving [NiFe]-enzymes, for example in *Escherichia coli* or *Wolinella succinogenes*^35,36^.

MR-1 is a biotechnologically significant organism and a central pillar of the study of [FeFe]-hydrogenase maturation. Its [FeFe]-hydrogenase belongs to a class from which only one enzyme has been described so far and it is found in an intriguing the metabolic context of C1 turnover - a task usually associated with [NiFe]-hydrogenase^35–38^. Therefore, the study of this enzyme contributes to better understanding of an important platform organism and could lead to better methods for its engineering. It will also shed new light on the diversity of [FeFe]-hydrogenases within enzyme families and highlight which features correlate with broad structural features and which ones are more strongly influenced by the fine-tuning of active site chemistry.

In this study we present the first biochemical characterisation of the [FeFe]-hydrogenase *So*HydAB, including activities, protein film electrochemistry, spectroscopy, and structural analysis. We show that this enzyme is structurally highly similar to *Nd*HydAB (Fig. 2), sharing some of its features but lacking others. The large and small subunits of the two enzymes share 49% and 33% sequence identity, respectively. The overall activity of *So*HydAB is much lower than that of *Nd*HydAB. Considering the consensus that hydrogenase activity is largely determined by small changes in direct proximity to the active site cofactor^39–41^, this is not as surprising as it may seem. One feature that stands out in the sequence of *So*HydA is an additional loop featuring two histidines and a tyrosine that are fully exposed.

**Figure 2:**
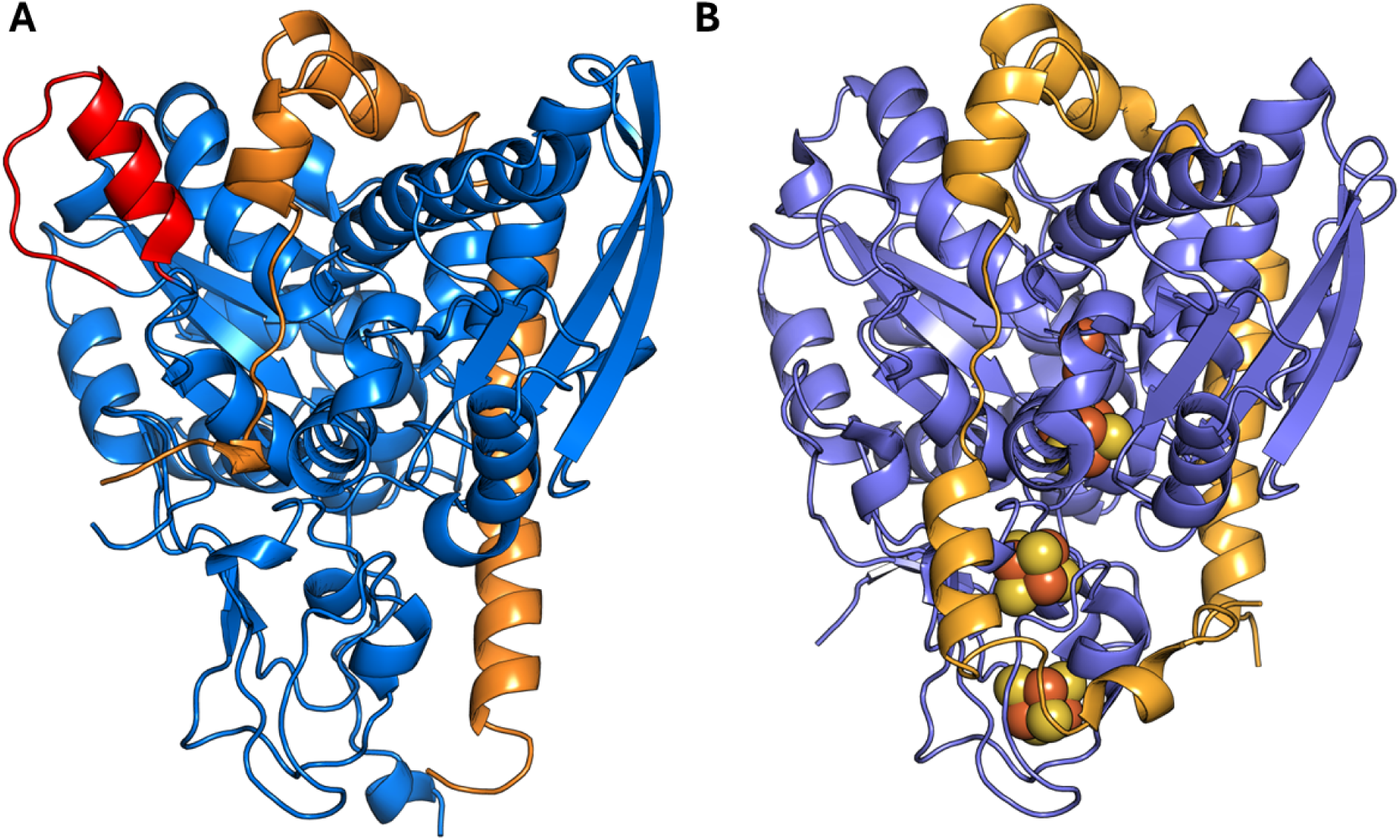
Side by side comparison of (A) *So*HyAB (AlphaFold3 model) and (B) *Nd*HydAB. (crystal structure PDB:1HFE^23^). Large subunits shown in blue, small subunits in orange. The extended loop in *So*HydAB is indicated in red. For overlay and further visual comparison see supplementary Fig. S1 and S2.

## Materials and Methods

### Protein Modelling

All predicted protein structures were generated using AlphaFold3^42^. All ribbon diagram figures were generated using PyMOL^43^.

### Enzyme Expression and Purification

Since the heterologous expression of HydA and HydB as separate proteins resulted in high losses of HydB and the cofactor, we designed a *So*HydAB fusion protein linked by glycine and serine residues (GS-linker). After evaluating constructs with four, six, eight and twelve residue linkers we settled on the GS-construct with six linkers (*So*HydA6B). Similar constructs for *Nd*HydAB have been shown to increase stability and cofactor integration without significantly affecting the catalytic behaviour of the enzyme^44^.

All genes were amplified from *S. oneidensis* MR-1 gDNA extracted using a Monarch Spin Plasmid Miniprep kit (New England Biolabs). Plasmids were constructed using NEBuilder HiFi DNA Assembly (New England Biolabs). Chemically competent cells of all utilised *E. coli* strains were prepared and transformed according to standard protocol. All variants of *So*HydAB were cloned in pET-vectors, amplified in *E. coli* NEB5alpha, and expressed in *E. coli* BL21 (DE3) Δ*iscR*^45^. For protein expression, cells were additionally transformed with pCYCHydEFXG, as presence of maturases was shown to boost hydrogenase yield.

Expression cultures were grown in LBP-medium (10 g/l peptone, 5 g/l yeast extract, 10 g/l Na2HPO4, 10 g/l K2HPO4) supplemented with antibiotics, 0.5% glucose and 2 mM ammonium iron citrate and inoculated with 3% main culture volume of preculture. After an OD_600_ of around 0.5 was reached, the culture was supplemented with 4 g/l sodium fumarate and moved to an anaerobic tent with 1.5–2.5% H_2_ atmosphere. After one hour of anaerobic growth, the culture was induced with 0.5mM Isopropyl β-D-1-thiogalactopyranoside (IPTG) and 0.6 g/l cysteine. Four hours later the cells were harvested (ten minutes at 6.000 rpm) and stored in airtight vessels at −20 °C.

For purification pellets were incubated in 100 mM Tris pH 8 (TPW) with 2 mg/l lysozyme and 1% Triton X-100 for 20 minutes and sonicated (five times 45 seconds with 1 minute breaks). The lysate was centrifuged for one hour at 40.000 rpm, filtered through a 0.2 µm membrane, applied to a Strep-Tactin-XT high-capacity or Strep-Tactin high-capacity column using an Äkta Start system and eluted with 50 mM biotin or 2.5 mM desthiobiotin, respectively. Elution fractions were pooled and concentrated using 30 K Amicon Ultra centrifugal filters (Millipore) and protein concentrations were determined via electronic spectroscopy at 260 nm. Size-exclusion chromatography (SEC) was carried out in SEC-buffer (10 mM Tris pH 8 and 100 mM NaCl) on a HiLoad 16/600 Superdex 200 pg column (Cytiva).

### Artificial Maturation

The H-cluster mimic was synthesised as previously described^46^. Enzymes were maturated by incubation at 18 °C for an hour with five-fold molar excess of the mimic. Excess mimic was removed by buffer exchange with TPW and reconcentrating in a 30 kDa centrifugal filter.

### Catalytic Activity

H_2_-reduction was measured via GC-TCD (GC-2010, Shimadzu) as described previously^47^. 2 ml of a solution containing 100 mM KP_i_ pH 6.8, 100 mM sodium dithionite (NaDT), 10 mM methyl viologen (MV), and 400 ng hydrogenase were prepared in a N_2_ atmosphere containing 1.5–2.5% H_2_ and sealed in an 8 ml glass vials using a rubber stopper. These samples were gassed with Argon for four minutes and incubated in a shaking water bath at 37 °C for 20 minutes. After incubation, 400 µl of the headspace gas were injected into the GC.

H_2_-oxidation activity was determined by electronic spectroscopy as previously described^48^. A cuvette was sealed with a rubber septum and sparged with 100% H_2_. 500 µl of a solution containing 100 mM KP_i_, 10 mM benzyl viologen (BV), and either 50 or 100 ng of hydrogenase were injected into the cuvette and the absorbance at 605 nm was measured over the course of two minutes. Activities were calculated based on the linear range of the absorbance function.

To determine the effects of various ions on activity, 500 mM of either KCl, NaCl, K-acetate, or Na-acetate were added to the reaction mixture.

### Electrochemistry

All protein film electrochemistry (PFE) experiments were carried out anaerobically. A PalmSens potentiostat (PalmSens 4) running PS trace 5.2 software was used to control the potential. The gas-tight electrochemical cell was water-jacketed to keep the temperature at 10 °C. A pyrolytic graphite “edge” rotating disk electrode controlled by a rotator (Autolab) was used as working electrode. A nonisothermal side arm housing the reference electrode (Ag/AgCl, 3.5 M KCl) was connected to the main cell compartment by a Luggin capillary. The reference potential was converted to standard hydrogen electrode (SHE), using an offset of −0.19 V. A platinum wire was used as a counter electrode. For each experiment, defined gas flow rates were set using a digital mass flow controller (Sierra Instruments).

For protein film preparation, 3 μl of 1 mg/ml hydrogenase solution was applied onto the polished electrode surface after measuring a blank. After 3 minutes, the electrode was rinsed with ultrapure water. All cyclic voltammetry (CV) experiments were performed at 10 °C and 3,000 rpm rotation frequency, under a 100% H_2_ atmosphere, using a scan rate of 0.01 V/s in a mixed buffer system (15 mM each of MES (2-(N-morpholino)ethanesulfonic acid), HEPES (4-(2-hydroxyethyl)-1-piperazineethanesulfonic acid), TAPS ([tris-(hydroxymethyl)methylamino]propanesulfonic acid), CHES (2-(Cyclohexylamino)-ethane-1-sulfonic acid), and sodium acetate, as well as 0.1 M NaSO_4_ unless specified otherwise) at pH 7. To determine the influence of NaCl, buffers without NaSO_4_ and with varying concentrations of NaCl were used.

### FTIR Spectroscopy

Experiments were conducted in an aerobic chamber (Coy Lab) utilising a FTIR spectrometer (Vertex 70, Bruker) equipped with a triple-reflection ZnSe/Si ATR optical cell. Infrared spectra were recorded at room temperature and a scanning velocity of 120 kHz with a spectral resolution of 2 cm^−1^ with a mercuric cadmium telluride (MCT) detector. The protein samples in the concentrations of 400 µM were applied on the ATR crystal, carefully dried, and rehydrated as described earlier^49^. To enrich H-cluster states, the sample films were flushed with changing composition of N_2_ and H_2_ at a flow rate of 1 l/min controlled by digital mass flow controllers (Sierra Instruments).

FTIR spectra in the CO/CN^−^ regime of the H-cluster were corrected for the background contribution of liquid H_2_O by the rubber band correction tool in OPUS (Bruker). Second derivative spectra of unmodified absorbance spectra were computed to verify the peak position of each band. Band fitting was performed using custom software tool GloFit. First, the CO/CN^−^ pattern of H_ox_ and H_ox_-CO was extracted from spectra showing an enrichment of these states including frequency, intensity, and FWHM. Spectra with mixed populations were described with H_ox_ and H_ox_-CO first before the remaining bands were assigned to the one- and two-electrons reduced species H_red_ and H_hyd_.

## Results

### Enzyme Expression and Purification

The expression of HydA and HydB in *E. coli* as separate proteins resulted in high losses of HydB and a lack of H-cluster integration. *So*HydA6B was expressed in *E. coli* BL21 DE3 Δ*iscR* with typical yields of around 25 mg of protein per litre of bacterial culture. The protein could be purified with high efficiency via strep tag. The purified enzyme could be concentrated up to 100 mg/ml (1.85mM) in TPW. On a size exclusion column the protein elutes as a monomer of 54 kDa. *So*HydA6B was fully maturated within an hour. This is a strong contrast to *Nd*HydAB which takes approximately 50 hours to fully mature this way^22^.

### Catalytic Activities

*So*HydA6B shows H_2_ evolution activities of 32 U/nmol (equivalent to 590 U/mg, MV) and H_2_ oxidation activities of 150 U/nmol (equivalent to 2860 U/mg, BV) under standard assay conditions at pH 6.8. We could show that the addition of Na^+^, K^+^, Cl^−^, or acetate-ions had no significant impact on activities of either *So*HydA6B or *Nd*HydAB (Fig. S7). This experiment was based on a 1979 study by Van Dijk *et al.*^50^, where the addition of 500 mM NaCl significantly boosted activities of the [FeFe]-hydrogenase from *Megasphaera*. This finding contrasts with the drastically altered activities of other [FeFe]-hydrogenases of M1, M2, and Mc types in the presence of these ions (Fig. S7). This implies that *Nd*HydAB and *So*HydA6B share characteristic properties probably related to surface charge.

### Electrochemistry

In PFE experiments *Nd*HydAB shows a strong bias toward H_2_ oxidation and high reversibility (Fig. 3A). In the presence of chloride ions, H_2_ oxidation is reversibly inactivated beginning at an overpotential of ∼400 mV vs. SHE^15,24^ (Fig. 3B). The characteristic shape of *Nd*HydAB voltammograms provided a salient starting point for electrochemical investigations of *So*HydA6B. Comparing the two enzymes, as well as model hydrogenase *Cr* HydA1 from green alga *Chlamydomonas reinhardtii* it becomes apparent that they are not affected by chloride in the same manner.

**Figure 3:**
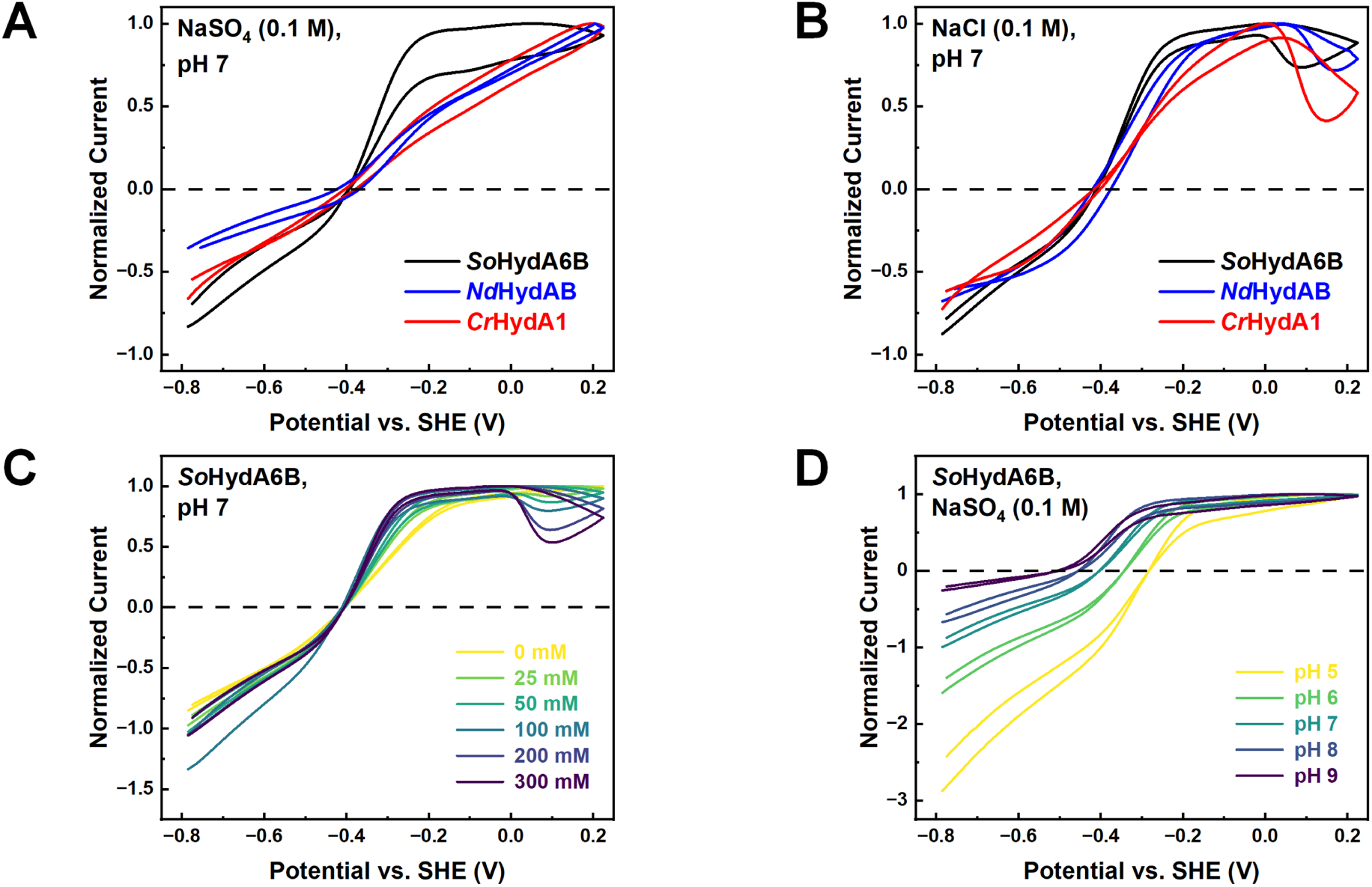
Electrochemical behaviour of *So*HydA6B and reference enzymes. All voltammograms recorded on a rotating graphite disk electrode, under H_2_ atmosphere in 15 mM mixed buffer pH 7. Measurements were taken at 10 °C and 3,000 rpm electrode rotation using a scan rate of 0.01 V/s. **(A)** Comparison of *So*HydA6B with *Nd*HydAB and *Cr* HydA1 with 0.1 M NaSO_4_. **(B)** Same experiment with 0.1 M NaCl instead of NaSO_4_. **(C)** *So*HydA6B with increasing concentrations of NaCl. **(D)** *So*HydA6B in mixed buffer with 0.1 M NaSO_4_ at different pH values. Currents normalised to maximal positive current.

*Cr* HydA1 exhibits an almost linear relationship between potential and current in the absence of chloride. With the addition of chloride there is a strong inactivation at positive potentials but up until that point reversibility is barely affected^51^. *Nd*HydAB behaves as outlined above, exhibiting an almost linear trace in the absence and increasingly more sigmoidal shape in the presence of chloride (Fig. 3C). In contrast, *So*HydA6B displays high currents for H_2_ evolution and H_2_ oxidation, with a slight catalytic bias toward H_2_ oxidation at pH 7 and extremely high reversibility with near-instantaneous switching between the reactions, rapidly reaching V_max_ for the H_2_ oxidation reaction. In the presence of chloride, the sigmoidal shape is even more pronounced and a slight inactivation at high potentials emerges. Comparing the behaviour of *So*HydA6B in the presence of different concentrations of NaCl (Fig. 3C) confirms the fast switching even in the complete absence of chloride (this is still exacerbated by chloride). Regardless of pH, the oxidative current of *So*HydA6B saturates at relatively low overpotentials (Fig. 3D).

### FTIR Spectroscopy

The “as isolated” *So*HydA6B sample shows an enrichment of the oxidized CO-inhibited H-cluster state H_ox_-CO (Fig. 4A). An unassigned band at 1983 cm^−1^ is observed. As described earlier^52^, the H_ox_-CO-to-H_ox_ conversion turned out to be a rather time-consuming effort. This process took at least two hours under continuous H_2_ flow (1 l/min) but was required for the enrichment of other H-cluster species. H_ox_ was accumulated to near-purity (Fig. 4A); however, the spectrum comprises an unassigned band at 1973 cm^−1^. Under 1% H_2_, H_ox_ quickly converted into a mixture of the one-electron reduced state H_red_ and the two-electrons reduced state H_hyd_ (Fig. 4A). The clear observation of H_hyd_ under 1% H_2_ and pH 6 is somewhat surprising as *Nd*HydAB has been shown to require pH 4 and 10% H_2_ for significant accumulation of Hhyd^53^. In general, fast reversible redox transitions between the H-cluster species were observed (Fig. 4B).

**Figure 4:**
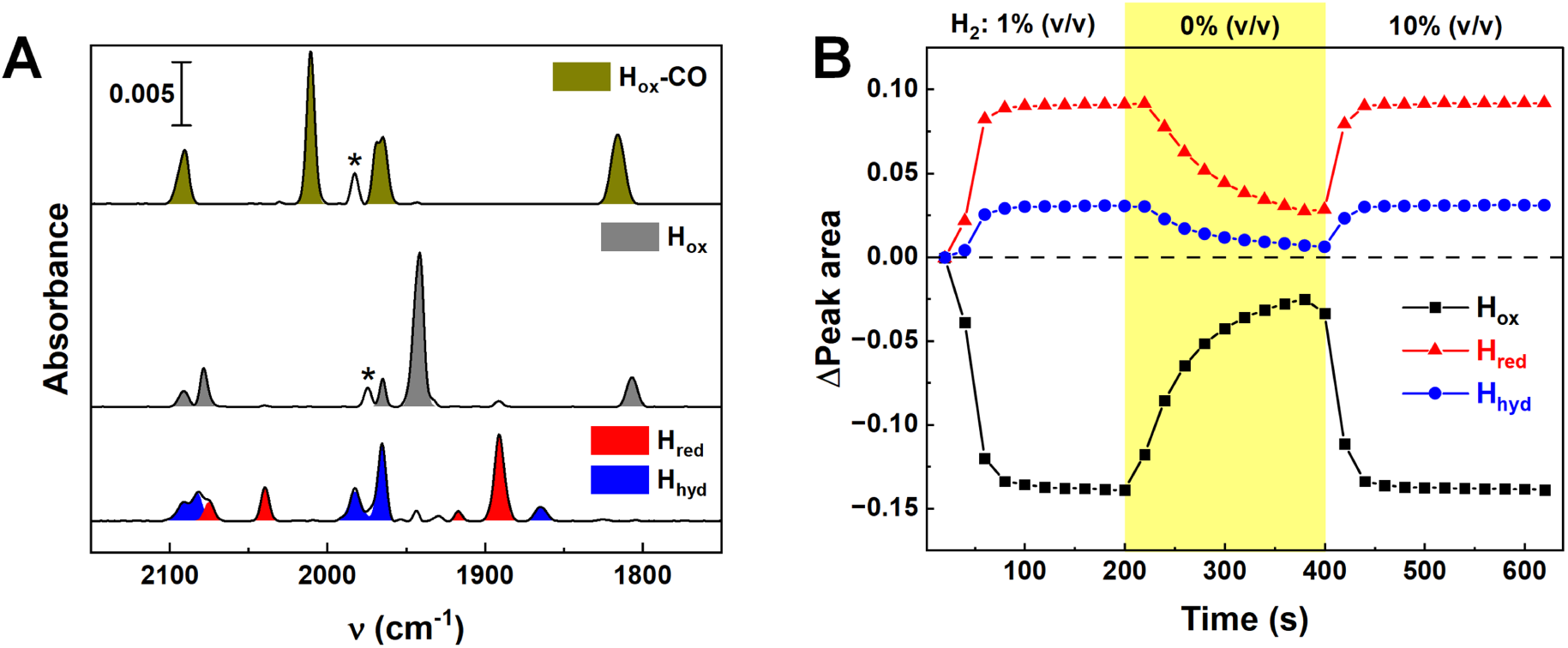
ATR FTIR spectroscopy of *So*HydA6B. **(A)** The “as isolated” sample shows an enrichment of the inhibited H_ox_-CO state (dark yellow). An unassigned band at 1983 cm^−1^ (*) is observed. After purging with N_2_ for 2–3 h, the H_ox_ state (grey) can be recovered. An unassigned band at 1973 cm^−1^ (*) is observed. In the presence of 1% H_2_, the H_red_ (red) and H_hyd_ (blue) states dominate. **(B)** The difference peak area of the catalytically active H-cluster states H_ox_, H_red_, and H_hyd_ (see Fig. S8) is plotted as a function of time. The data show how the H-cluster becomes fully reduced under 1% H_2_ and quickly “auto-oxidizes” toward H_ox_ under 100% N_2_ (yellow area). The ratio of reduced state H_red_ and H_hyd_ is the same under 1% and 10% H_2_.

After exposure of active, oxidized *So*HydA6B to O_2_, the H-cluster is deactivated within seconds: the H_ox_ signatures disappears, as does the unassigned band at 1973 cm^−1^ (Fig. S9). A very small fraction of H_ox_-CO becomes visible (positive band at 2010 cm^−1^) providing hints for the destruction of H-cluster species releasing CO that binds to other H-cluster species^54^. The assigned frequencies for each H-cluster species are given in Table 1.

**Table 1:** CO/CN frequencies (cm^−1^) and their assigned H-cluster states in *So*HydAB.

| | n(CN<sup>-</sup>) | n(CN<sup>-</sup>) | n(COp) | n(COd) | n(μCO) |
| --- | --- | --- | --- | --- | --- |
| H<sub>ox</sub>-CO | 2094 | 2086 | 1970 | 2010/1964 | 1816 |
| H<sub>ox</sub> | 2091 | 2079 | 1974 | 1941 | 1807 |
| H<sub>red</sub> | 2074 | 2039 | 1917 | 1891 | 1825 |
| H<sub>hyd</sub> | 2092 | 2082 | 1983 | 1965 | 1865 |

## Discussion

Despite the structural similarities, *So*HydAB does not behave quite like *Nd*HydAB. There are obvious differences in stability and the duration of artificial maturation. Unfused *So*HydAB seems to be less stable when purified from *E. coli*. One possible reason for this could be that *So*HydB is 17 amino acids shorter than its analogue in *Nitratidesulfovibrio* and does not wrap around the large subunit as fully (Fig. 1B). This decreased stability of the dimer could in turn allow for faster maturation. In fact, Winkler *et al.* recently reported efficient maturation in *Nd*HydAB variants related to HydB^44^.

The quick saturation of H_2_ oxidation activity at relatively low overpotential exhibited by *So*HydAB is an indication for the formation of an unusually orderly protein film on the electrode in which electron transport is not limiting activity. One could speculate that the interface that enables interaction with HydC (see below) is likewise ideally suited for interacting with an electrode surface but confirming this will require further investigation.

Besides these differences, *So*HydAB and *Nd*HydAB show many similarities – like being weakly impacted by ions (chloride, protons) and having similar electrochemical behaviour. The strength of the response, however, differs strongly between the enzymes, with *So*HydAB consistently exhibiting lower activities than *Nd*HydAB. As FTIR spectroscopy indicates a strong CO-inhibition of as-isolated sample, we assume that activities and PFE response underestimate the efficiency of the enzyme. The main determinant of the activity is the interplay of the microenvironments of the active site and the substrate channels – the proton transfer pathway, electron transport chain, and gas channels^41^. Similar trends have been observed between two isoforms of [FeFe]-hydrogenase in *C. reinhardtii* (*Cr* HydA1 and *Cr* HydA2) that share an overall shape and exhibit similar electrochemical behaviour while differing drastically in activity^55^. We propose that this could be an adaptation to the different electron sources, with *Nd*HydAB “competing” for reduced cytochrome c with other potential electron acceptors^56^ while *So*HydAB is supplied with electrons directly by HydC^11^.

The interaction between HydAB and HydC is thermodynamically plausible based on AlphaFold3 models (Fig. S1) but has not been explored in experiments. The electron mediator MQ that facilitates electron transfer between FDH, HydC, and HydAB in *S. oneidensis* appears to be specific, given that the deletion of hydC abolishes H_2_ production by HydAB^11^. We speculate that the interaction of HydAB and HydC could be dynamic and redox-depend, similar to the electron transfer complex of [FeFe]-hydrogenase and ferredoxin^57^. However, upon reduction of *So*HydA6B alone, FTIR difference spectra in the amide regime (Fig. S8 and Fig. S10) do not suggest protein structural change that could modulate a dynamic interaction with HydC. This might be different in presence of HydC, and respective experiments are currently planned. *So*HydA6B being rendered less flexible than the native dimer by the GS-linker is another possibility.

*Shewanella oneidensis* has been known to oxidise H_2_ under certain conditions, however, the role [FeFe]-hydrogenases play in this has been discussed controversially^9,58,59^. We have shown now that *So*HydAB is clearly capable of bidirectional catalysis *in vitro*. Given that the expression of hydrogenases is associated with acidification, HydAB is likely biased towards proton reduction (H_2_ evolution) under physiological conditions, which agrees with our electrochemical results. We propose that the connection to the menaquinone pool facilitates a constant influx of electrons while H_2_ in the periplasm either diffuses out of the cell or is taken up by the [NiFe]-hydrogenase HyaAB, due to its higher H_2_ affinity. This is consistent with previous results showing no physiological H_2_ oxidation by HydAB^9^. We note that *So*HydAB readily accumulates the two-electrons reduced H_hyd_ state *in vitro*, indicating good compatibility with MQH_2_, a two-electron donor.

Xiong *et al.*^11^ deduced from the necessity of menaquinone that *S. oneidensis* does not have a formate-hydrogen lyase (FHL) complex. We consider this an overinterpretation, given that a reliance on menaquinone does not rule out a more structured interaction between FdH and the hydrogenase. If that interaction is similar enough to a canonical FHL to deserve that designation is a matter of further research and debate. We merely want to caution that it is too early to rule out an FHL-like complex based on current data.

This study shows that even structurally highly similar [FeFe]-hydrogenases can exhibit considerable variation in their biochemical properties and showcases the diversity of [FeFe]-hydrogenases not only between but also within enzyme families. This contributes to a better understanding of which properties can actually be predicted by looking at phylogeny or structure. Furthermore, we want to foster renewed interest in hydrogen metabolism in *Shewanella oneidensis*, particularly on a molecular level. The strategy presented here should provide a foundation for further investigations into HydC and the concrete mechanisms of electron transfer from formate to protons in *Shewanella oneidensis*.

## Acknowledgements

We thank Prof. Ulf-Peter Apfel and Dr. Julian Kleinhaus for providing the H-cluster mimic. We thank Jan Jaennecke for providing *Nd*HydAB. We acknowledge discussing formate turnover with Prof. Gary Sawers.

This research was funded by the DFG Research Training Group GRK 2341 “Microbial Substrate Conversion” (MiCon). We thank the complete Research Training Group GRK 2341 “MiCon” for support of our graduate students. S.T.S. acknowledges the German Research Foundation for funding (DFG grants STR 1554/6-1 and STR 1554/8-1).

## Supplementary Information

**Supplementary Figure S 1:**
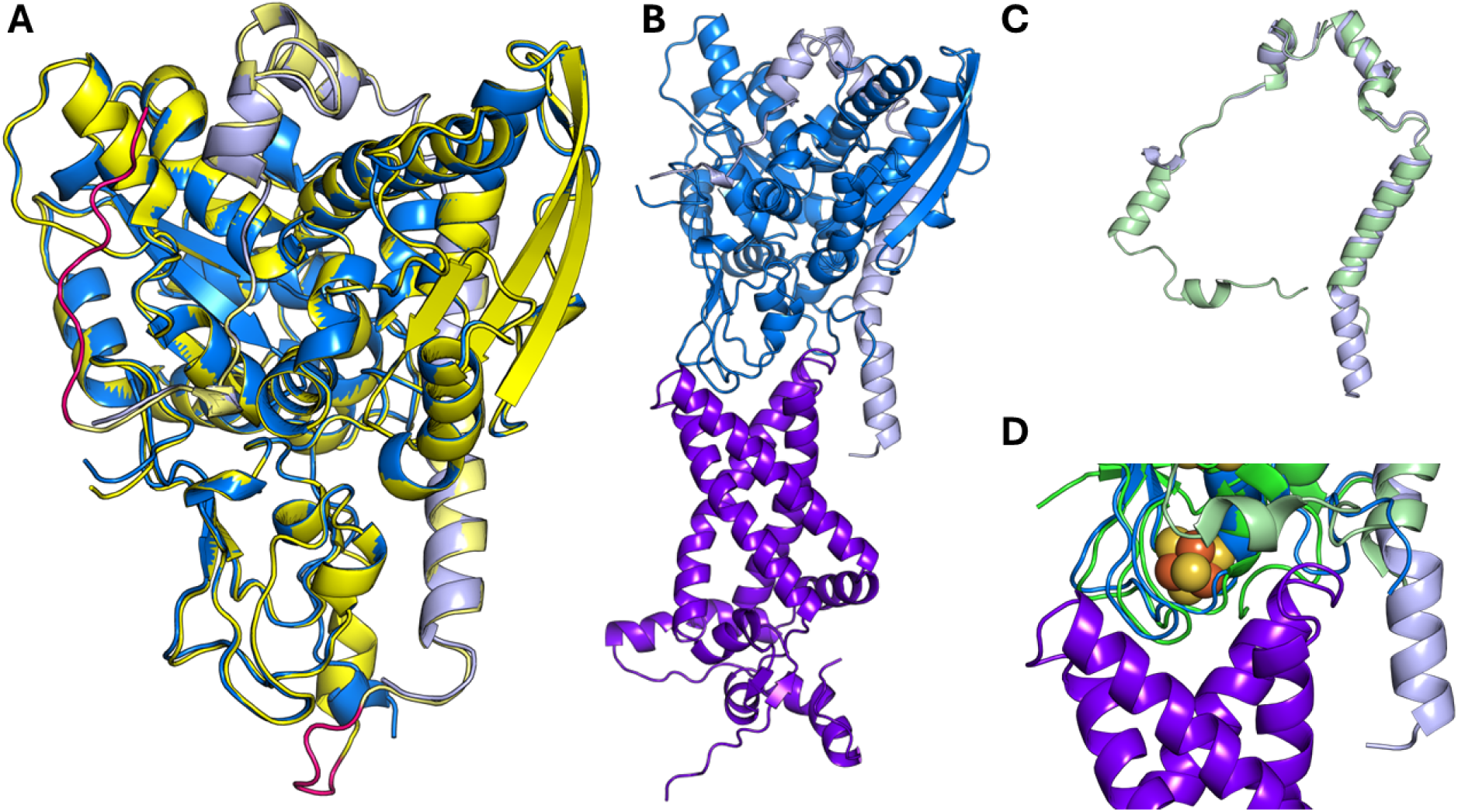
AlphaFold 3 models of *So*HydABC (HydA shown in blue, HydB in light blue, HydC in purple) compared to the crystal structure of *Nd*HydAB (1hfe, HydA in green, HydB in light green). **(A)** Alignment of native *So*HydAB with *So*HydA6B. Fusion protein in yellow, linker and strep-tag highlighted in red. **(B)** HydABC. HydC shows a canonical cytochrome b fold. Notice the short N-terminal helix on HydB (IPIGWFTSKLE) likely entering into the membrane. **(C)** Alignment of the small subunits (*So*HydB in blue, *Nd* HydB in green). In *Nd*HydAB the small subunit completely wraps around the large subunit but cannot extend a helix away from the protein core. **(D)** Alignment of *So*HydABC with *Nd*HydAB (FeS cluster from crystal structure), highlighting the difference in the region around the distal F-cluster.

**Supplementary Figure S 2:**
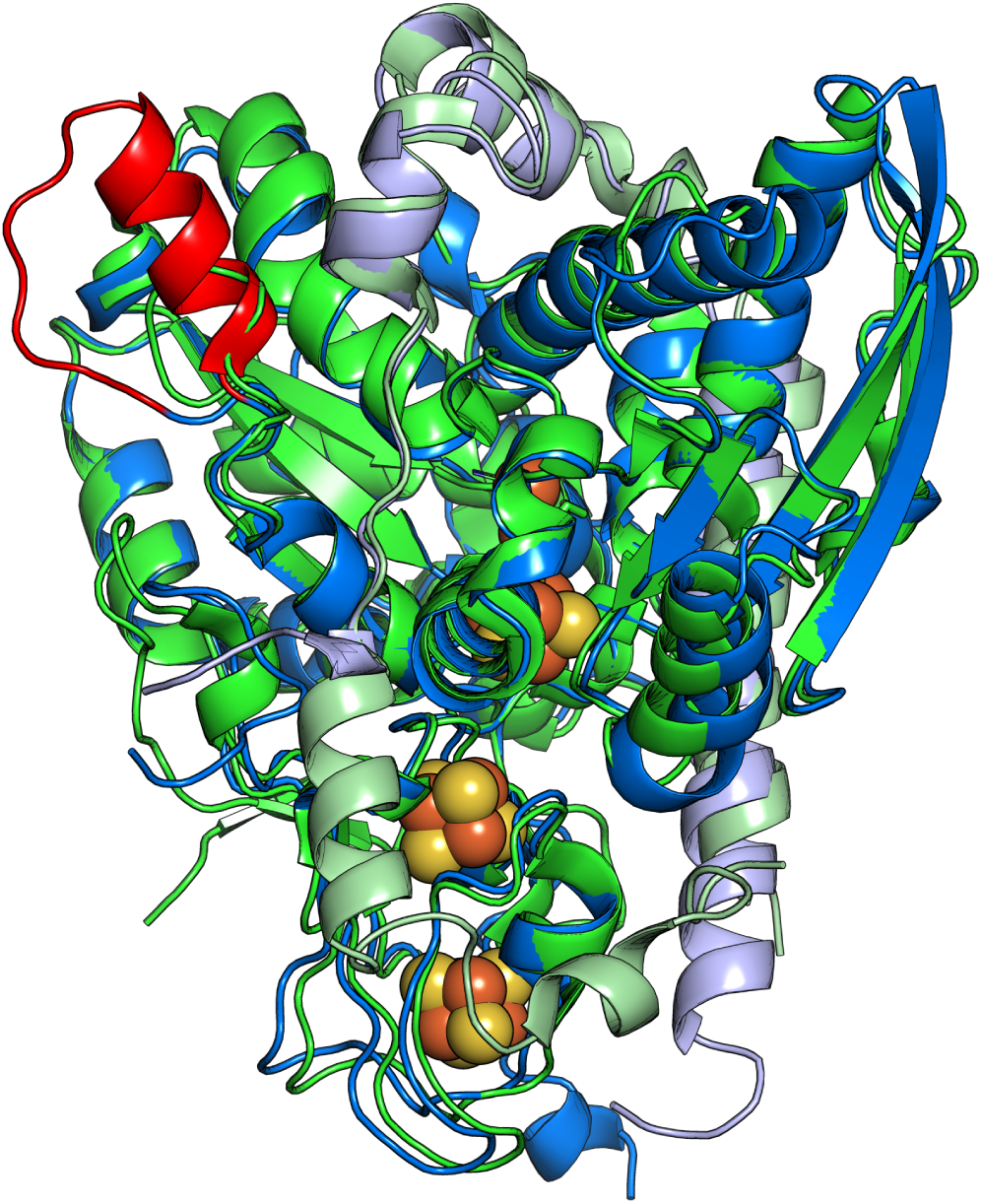
Structural Alignment of *Nd*HydAB (1hfe, HydA in green, HydB light green) and an AlphaFold 3-model of *So*HydAB (HydA blue, HydB light blue, inserted loop red).

**Supplementary Figure S 3:**
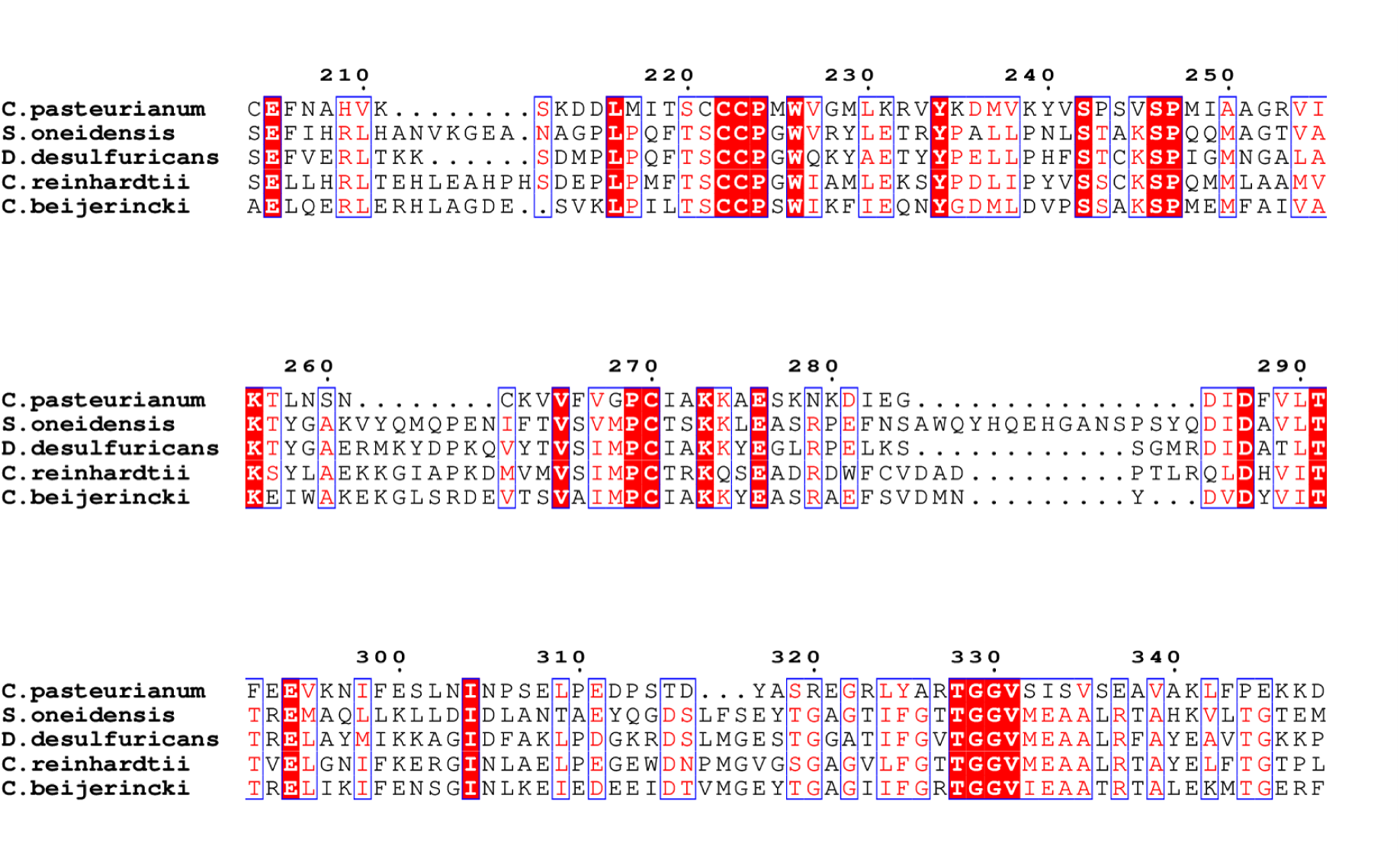
Amino acid sequence alignment of *So*HydAB with model [FeFe]-hydrogenases. (*C. pasteurianum* - *Cp*I, *N. desulfuricans* - *Nd* HydA, *C. reinhardtii* - *Cr* HydA1 and *C. beijerincki Cb*A5H). Note the loop shown in main article figure 3 that is absent in all other enzymes.

**Supplementary Figure S 4:**
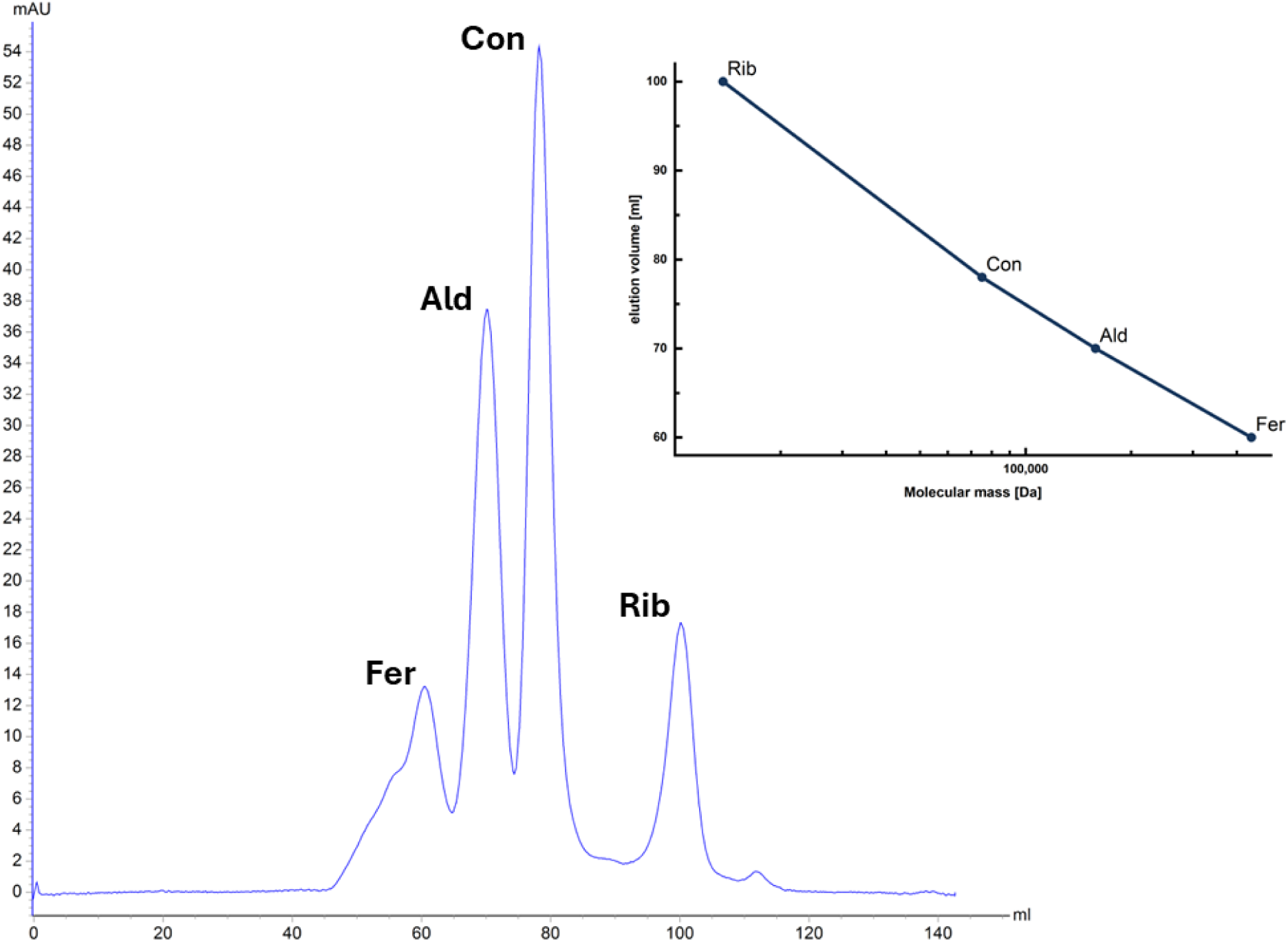
Chromatogram of the Size Exclusion Chromatography (SEC) Calibration. Chromatography was carried out in SEC-buffer (10mM Tris pH 8 100mM NaCl) on a HiLoad 16/600 Superdex 200pg column (Cytiva). The proteins used for calibration and their molecular masses (in Da) are Ferritin (440 000), Aldolase (158 000), Conalbumin (75 000) and Ribonuclease A (13 700). Inset shows a linear relationship between molar mass and elution volume, allowing reliable estimation of protein mass.

**Supplementary Figure S 5:**
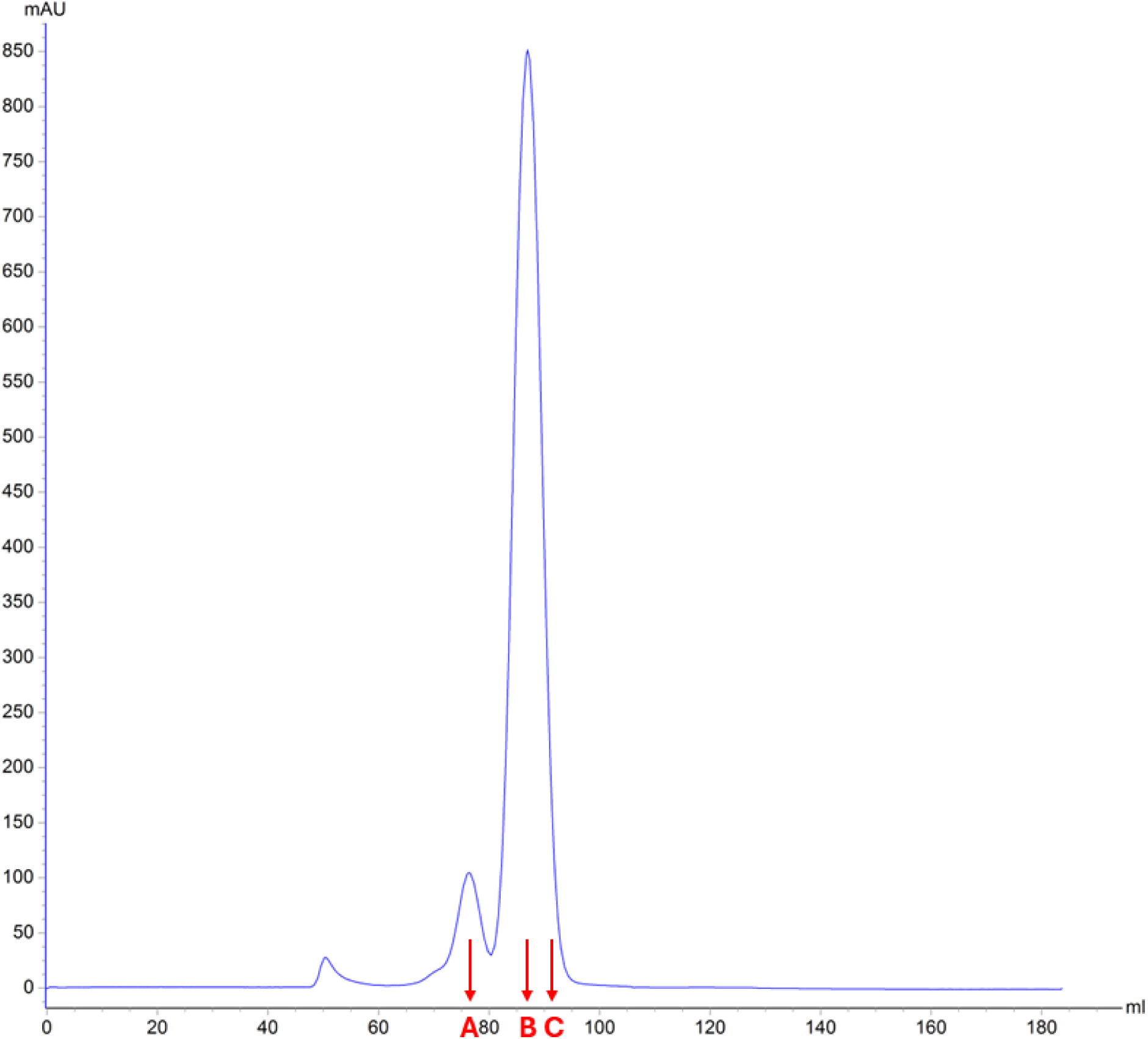
Size Exclusion chromatogram of *So*HydA6B. SEC was carried out in SEC-buffer (10mM Tris pH 8 100mM NaCl) on a HiLoad 16/600 Superdex 200pg column (Cytiva). Samples from points A, B and C can be seen on SDS-PAGE in figure S5.

**Supplementary Figure S 6:**
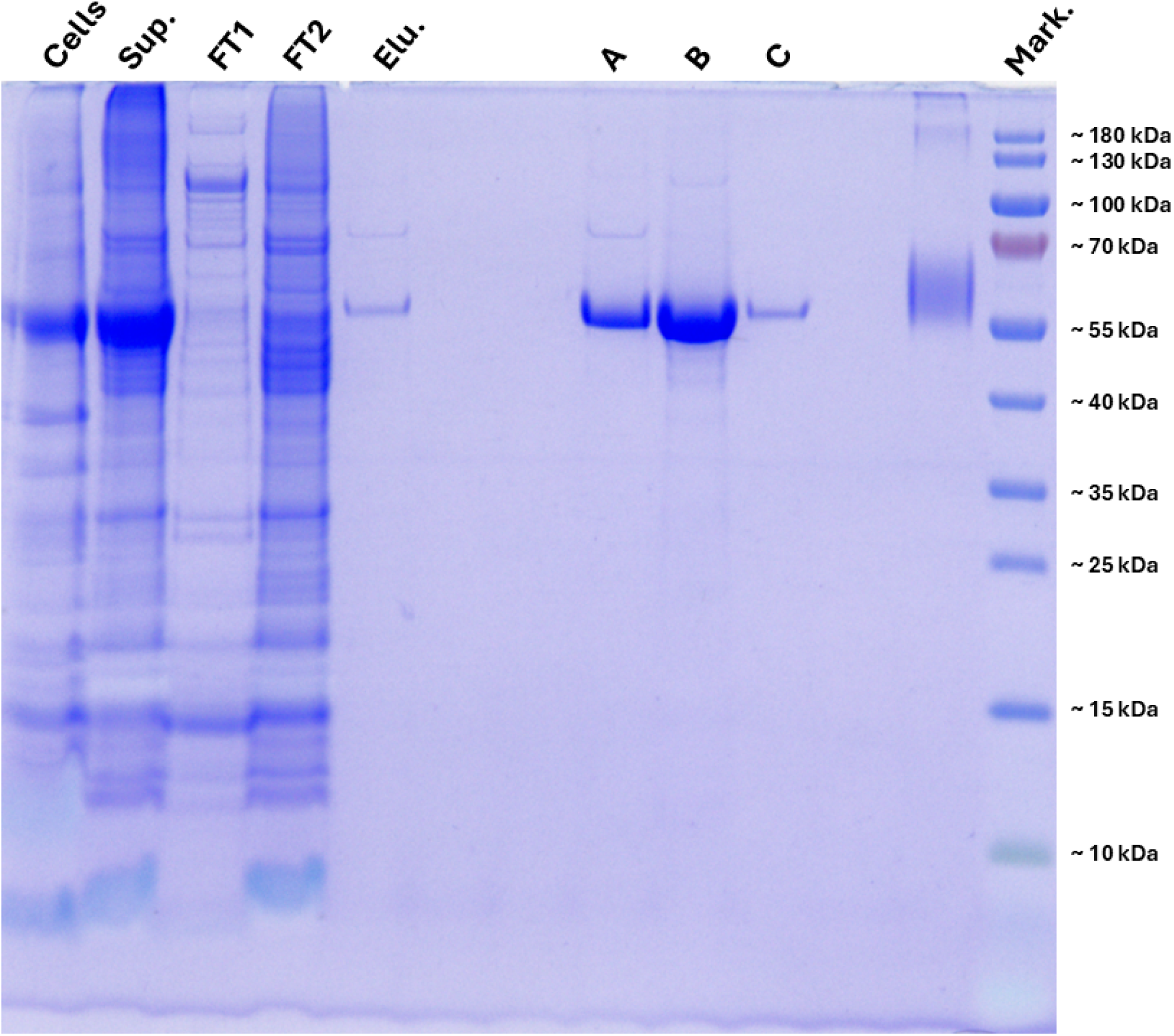
SDS-PAGE gel of purified *So*HydA6B. Cells: Expression culture (*E. coli* BL21 (DE3) Δ*iscR* expressing *So*HydA6B, see Methods section) at time of harvest. Sup.: Supernatant after sonication and centrifugation. FT1: Early flow-through on Strep-Tactin-XT high capacity column. FT2: late flow-through. Elu: pooled elution fractions. A, B, C: gel filtration samples highlighted in figure S4. Marker: Thermo Scientific™ PageRuler Prestained Protein Ladder. Samples were loaded onto a 12% Bis-Tris gel and electrophoresis was performed at 200V, 120mA for 40 minutes in SDS-MOPS buffer.

**Supplementary Figure S 7:**
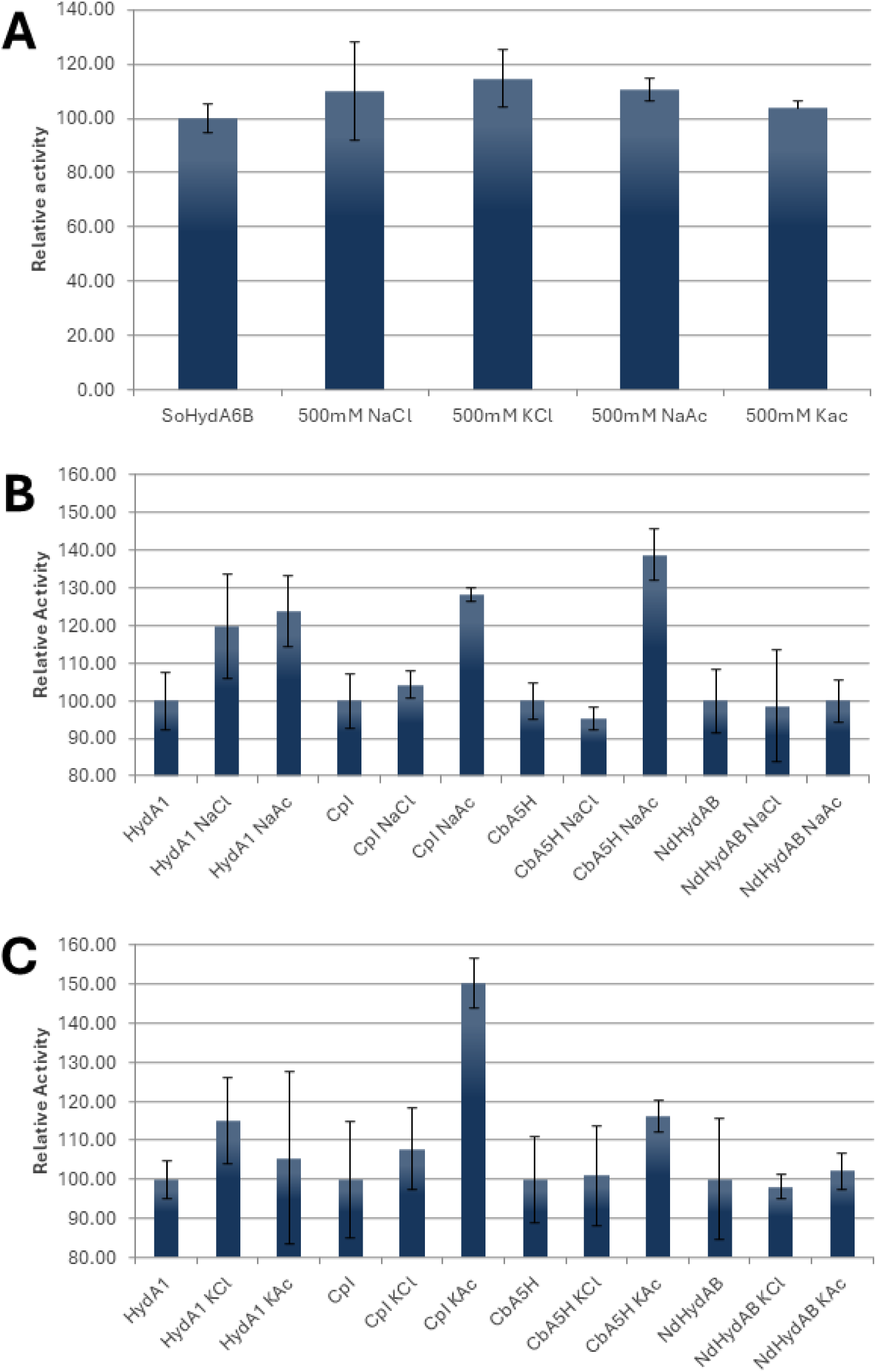
Ion effects on activities of various [FeFe]-hydrogenases. All measurements were carried out on 400µl of headspace gas from a reaction mixture containing 100mM KP_i_ pH 6.8, 100mM NaDT, 10mM methyl viologen and 400 ng of the respective hydrogenase. **(A)** addition of the indicated salt or no additional salts. **(B)** addition of 500mM NaCl, NaAc or no additional salts. **(C)** addition of 500mM KCl, KAc or no additional salts.

**Supplementary Figure S 8:**
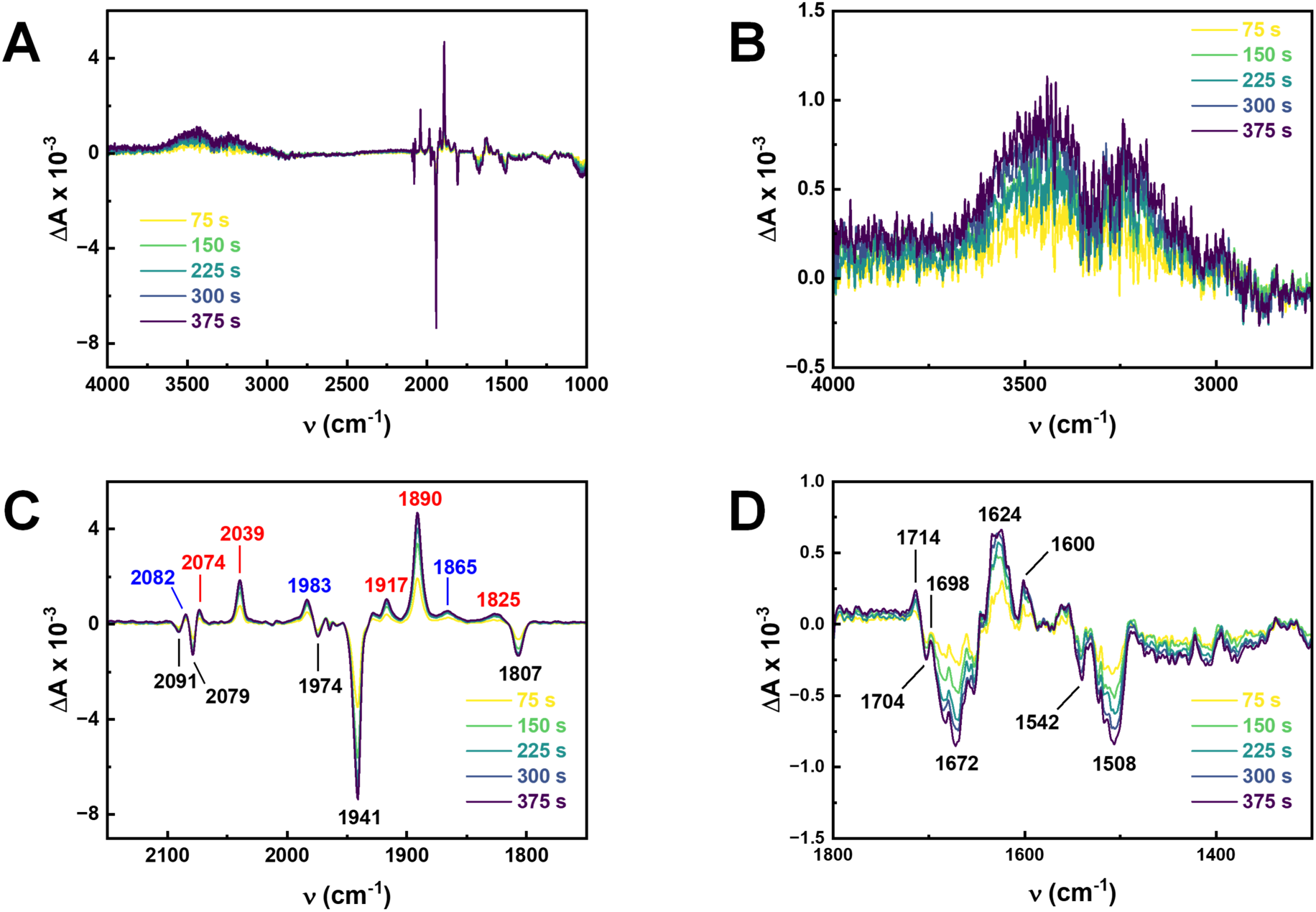
Time series of H_2_–N_2_ FTIR difference spectra that track the reduction of *So*HydAB. **(A)** Overview spectrum between 4000–1000 cm^−1^. Note the stable baseline and lack of “unspecific” changes in hydration or protein concentration. **(B)** Upon closer inspection, the spectra hint at a minor, negligible increase of hydration at higher frequencies, marked by the OH oscillators of H_2_O. **(C)** The CO/CN regime of the H-cluster reveals the accumulation of positive bands assigned to H_red_ (red) and H_hyd_ (blue) over the negative bands of H_ox_. **(D)** At lower frequencies, bands related to protonation changes in the PTP become visible (1714, 1704, and 1698 cm^−1^). Small signals in the Amide I regime (1672, 1624, and 1600 cm^−1^) and Amide II regime (1542 and 1508 cm^−1^) suggest very small protein structural changes. See Fig. S10 for a clear detection of protein structural changes in another [FeFe]-hydrogenase.

**Supplementary Figure S 9:**
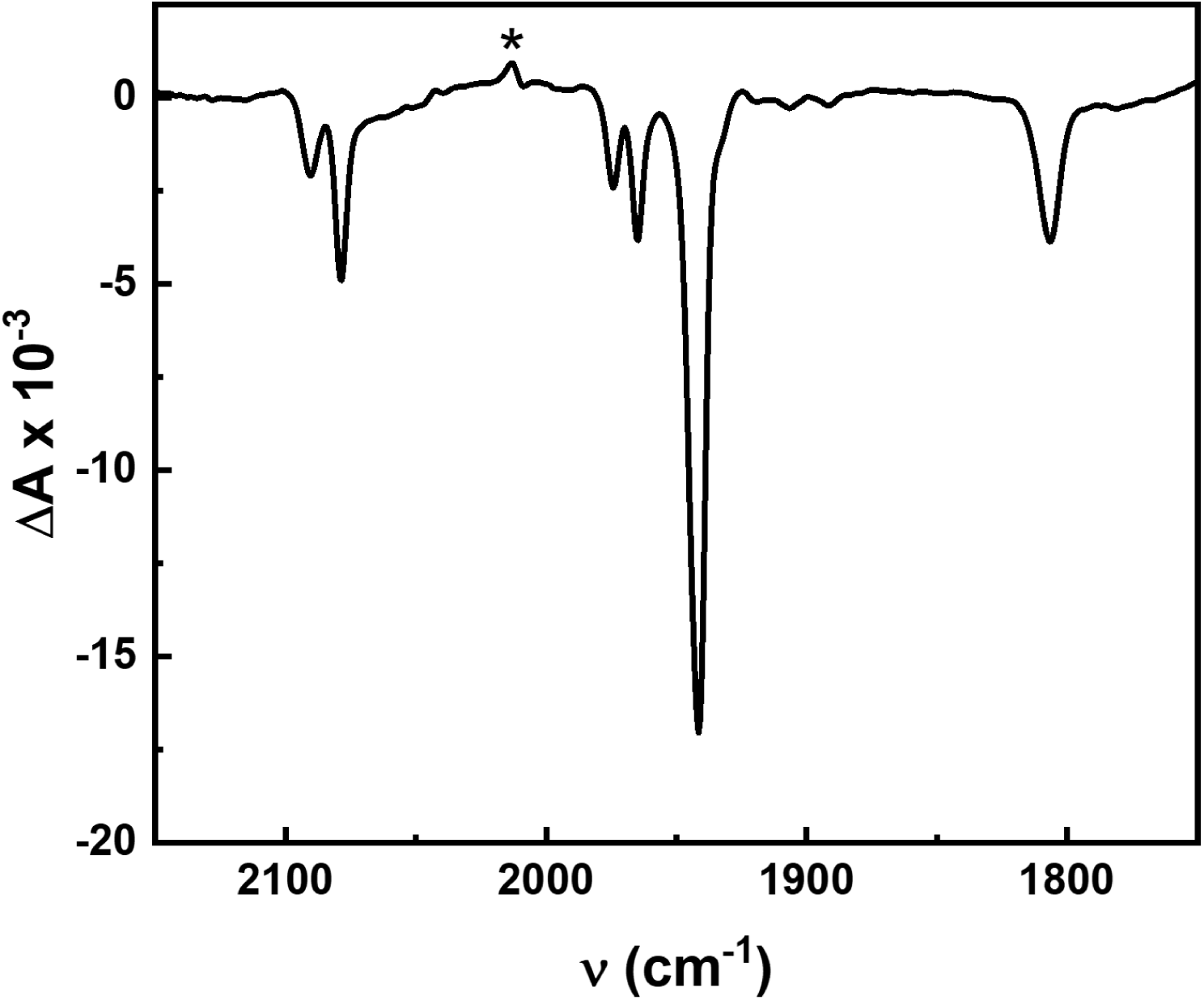
O_2_–N_2_ FTIR difference spectrum. After activation of the “as isolated” sample under N_2_ for 2–3 h toward full accumulation of the H_ox_ state, SoHydA6B was reacted with 1 atm O_2_, which led to an immediate loss of all cofactor bands. Calculating the O_2_–N_2_ difference spectrum reveals a small fraction of H_ox_-CO (*) as the result of cannibalization.

**Supplementary Figure S 10:**
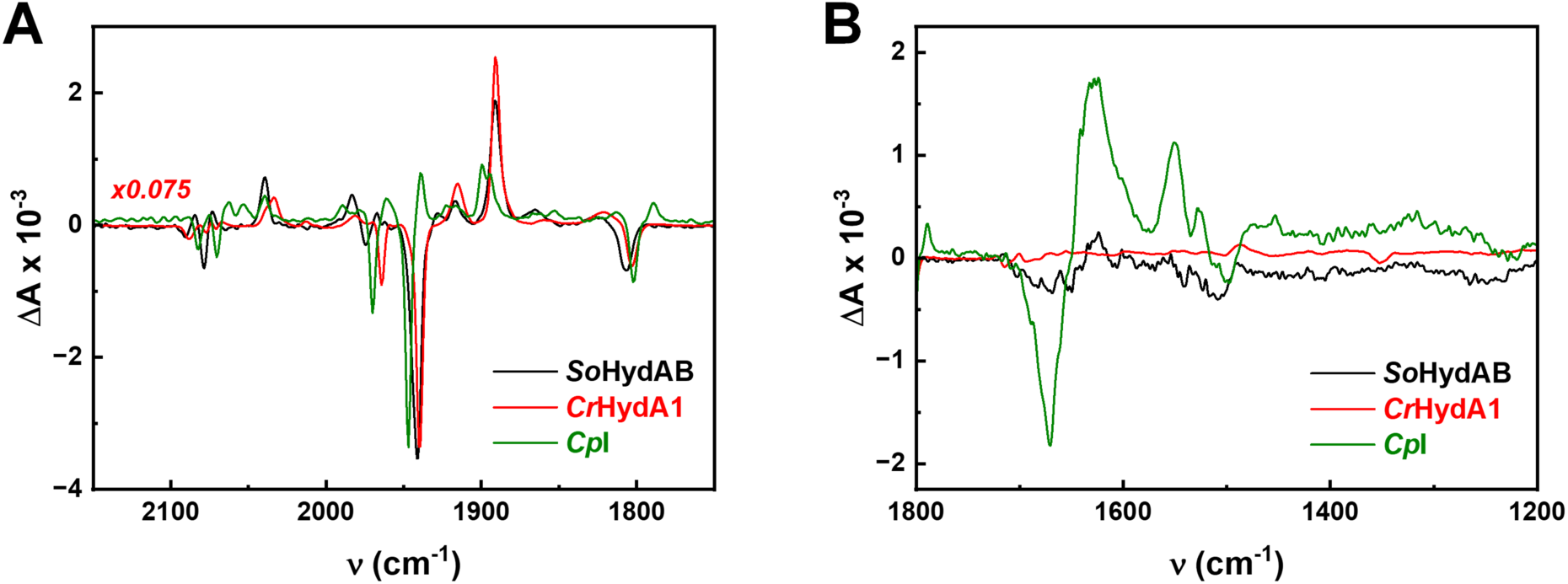
Comparison of H_2_–N_2_ FTIR difference spectra of different [FeFe]-hydrogenases. **(A)** In the CO/CN regime of the H-cluster, accumulation of H_red_ over H_ox_ is observed for *So*HydAB and *Cr* HydA1. In *CpI* of *Clostridium pasteurianum*, H_red_ is accumulated over H_ox_ alongside H_sred_ and H_red′_. **(B)** At lower frequencies, *Cp*I shows clear Amide I difference bands that are related to changed in the F-domain. *Cr* HydA1, which lacks the F-domain, shows no such signals. In comparison to *Cp*I, the Amide-related difference signals in *So*HydAB (Fig. S8) are very small and cannot explain a redox-dependent interaction with HydC.

## Amino acid sequence of the *So*HydA6B construct used in this study

M T T T T Y Q P G E I Q G L I K I N A S K C K G C D A C K Q F C P T H A I N G A S G A V H S I D E D K C L S C G Q C L I N C P F S A I E E T H S A L E T V I K K L A D K N T T V V G I I A P A V R V A I G E E F G L G T G E L V T G K L Y G A M N Q A G F K I F D C N F A A D L T I M E E G S E F I H R L H A N V K G E A N A G P L P Q F T S C C P G W V R Y L E T R Y P A L L P N L S T A K S P Q Q M A G T V A K T Y G A K V Y Q M Q P E N I F T V S V M P C T S K K L E A S R P E F N S A W Q Y H Q E H G A N S P S Y Q D I D A V L T T R E M A Q L L K L L D I D L A N T A E Y Q G D S L F S E Y T G A G T I F G T T G G V M E A A L R T A H K V L T G T E M A K L E F E P V R G L K G V K S A S V S L F D T E L N Q D V T V N V A V V H D M G N N I E P V L R D V M A G T S P Y H F I E V M N C A G G C V N G G G Q P I E G K G S S W L G N I S G G S G S I P I G W F T S K L E R R N E Y I K A R S Q G L Y K D D S L A K T R V S H A N P A V E K Y Y K E F G G E P L G H M S H E L L H T H F V D R T K L S S S A W S H P Q F E K

## References

1. Venkateswaran, K. et al. Polyphasic taxonomy of the genus *Shewanella* and description of *Shewanella oneidensis* sp. nov. Int. J. Syst. Bacteriol. 49, 705–724 (1999).

2. Myers, C. R., Carstens, B. P., Antholine, W. E. & Myers, J. M. Chromium(VI) reductase activity is associated with the cytoplasmic membrane of anaerobically grown *Shewanella putrefaciens* MR-1. J. Appl. Microbiol. 88, 98–106 (2000).

3. Windt, W. De, Aelterman, P. & Verstraete, W. Bioreductive deposition of palladium (0) nanoparticles on *Shewanella oneidensis* with catalytic activity towards reductive dechlorination of polychlorinated biphenyls. Environ. Microbiol. 7, 314–325 (2005).

4. Fredrickson, J. K., et al. Reduction of U(VI) in Goethite (-FeOOH) Suspensions by a Dissimilatory Metal-Reducing Bacterium. (2000).

5. Hau, H. H. & Gralnick, J. A. Ecology and biotechnology of the genus *Shewanella*. Annual Review of Microbiology vol. 61 237–258 Preprint at 10.1146/annurev.micro.61.080706.093257 (2007).

6. Pirbadian, S., et al. *Shewanella oneidensis* MR-1 nanowires are outer membrane and periplasmic extensions of the extracellular electron transport components. Proc. Natl. Acad. Sci. U. S. A. 111, 12883–12888 (2014).

7. Dianne K. Newman & Roberto Kolter. A role for excreted quinones in extracellular electron transfer. Letters to nature (2000).

8. Beliaev, A. S. et al. Global transcriptome analysis of *Shewanella oneidensis* MR-1 exposed to different terminal electron acceptors. J. Bacteriol. 187, 7138–7145 (2005).

9. Meshulam-Simon, G., Behrens, S., Choo, A. D. & Spormann, A. M. Hydrogen metabolism in *Shewanella oneidensis* MR-1. Appl. Environ. Microbiol. 73, 1153–1165 (2007).

10. Kneuer, L., Wurst, R. & Gescher, J. *Shewanella oneidensis*: Biotechnological Application of Metal-Reducing Bacteria. Adv. Biochem. Eng. Biotechnol. 192, 117–154 (2025).

11. Xiong, J. et al. Hydrogen production driven by formate oxidation in *Shewanella oneidensis* MR-1. Appl. Microbiol. Biotechnol. 104, 5579–5591 (2020).

12. Agapakis, C. M., et al. Insulation of a synthetic hydrogen metabolism circuit in bacteria. J. Biol. Eng. http://www.jbioleng.org/content/4/1/3 (2010).

13. Gärtner, K., Lechno-Yossef, S., Cornish, A. J., Wolk, C. P. & Hegg, E. L. Expression of *Shewanella oneidensis* MR-1 [FeFe]-hydrogenase genes in *Anabaena* sp. strain PCC 7120. Appl. Environ. Microbiol. 78, 8579–8586 (2012).

14. Lubitz, W., Ogata, H., Rüdiger, O. & Reijerse, E. Hydrogenases. Chem. Rev. 114, 4081–4148 (2014).

15. Vincent, K. A., Parkin, A. & Armstrong, F. A. Investigating and exploiting the electrocatalytic properties of hydrogenases. Chem. Rev. 107, 4366–4413 (2007).

16. Mulder, D. W. et al. Insights into [FeFe]-Hydrogenase Structure, Mechanism, and Maturation. Structure Review 10.1016/j.str.2011.06.008 (2011) doi:10.1016/j.str.2011.06.008.

17. Kuchenreuther, J. M., Britt, R. D. & Swartz, J. R. New Insights into [FeFe] Hydrogenase Activation and Maturase Function. PLoS One 7, (2012).

18. Lampret, O. et al. The final steps of [FeFe]-hydrogenase maturation. Proc. Natl. Acad. Sci. U. S. A. 116, 15802–15810 (2019).

19. Sybirna, K., et al. *Shewanella oneidensis*: A new and efficient System for Expression and Maturation of heterologous [Fe-Fe] Hydrogenase from *Chlamydomonas reinhardtii*. BMC Biotechnol. 8, (2008).

20. Meyer, J. [FeFe] hydrogenases and their evolution: a genomic perspective. Cellular and Molecular Life Sciences 2007 64:9 64, 1063–1084 (2007).

21. Land, H., Senger, M., Berggren, G. & Stripp, S. T. Current State of [FeFe]-Hydrogenase Research: Biodiversity and Spectroscopic Investigations. ACS Catalysis vol. 10 7069–7086 Preprint at 10.1021/acscatal.0c01614 (2020).

22. Birrell, J. A. et al. Artificial Maturation of the Highly Active Heterodimeric [FeFe] Hydrogenase from *Desulfovibrio desulfuricans* ATCC 7757. Isr. J. Chem. 56, 852–863 (2016).

23. Nicolet, Y., Piras, C., Legrand, P., Hatchikian, C. E. & Fontecilla-Camps, J. C. Desulfovibrio desulfuricans iron hydrogenase: the structure shows unusual coordination to an active site Fe binuclear center. Structure http://biomednet.com/elecref/0969212600700013 (1999).

24. Parkin, A., Cavazza, C., Fontecilla-Camps, J. C. & Armstrong, F. A. Electrochemical investigations of the interconversions between catalytic and inhibited states of the [FeFe]-hydrogenase from *Desulfovibrio desulfuricans*. J. Am. Chem. Soc. 128, 16808–16815 (2006).

25. Yagi, T., Honya, M. & Tamiya, N. Purification and properties of hydrogenases of different origins. Biochimica et Biophysica Acta (BBA) - Bioenergetics 153, 699–705 (1968).

26. Heidelberg, J. F. et al. Genome sequence of the dissimilatory metal ion-reducing bacterium *Shewanella oneidensis*. Nat. Biotechnol. 20, 1118–1123 (2002).

27. Gross, R., Pisa, R., Sänger, M., Lancaster, C. R. D. & Simon, J. Characterization of the Menaquinone Reduction Site in the Diheme Cytochrome b Membrane Anchor of *Wolinella succinogenes* NiFe-hydrogenase. Journal of Biological Chemistry 279, 274–281 (2004).

28. Kane, A. L. et al. Formate metabolism in *Shewanella oneidensis* generates proton motive force and prevents growth without an electron acceptor. J. Bacteriol. 198, 1337–1346 (2016).

29. Pinchuk, G. E. et al. Pyruvate and lactate metabolism by *Shewanella oneidensis* MR-1 under fermentation, oxygen limitation, and fumarate respiration conditions. Appl. Environ. Microbiol. 77, 8234–8240 (2011).

30. Brosnan, M. E. & Brosnan, J. T. Formate: The Neglected Member of One-Carbon Metabolism. Annu. Rev. Nutr. 36, 369–388 (2016).

31. Lü, W. et al. pH-dependent gating in a FocA formate channel. Science (1979). 332, 352–354 (2011).

32. Lo, Y. K. et al. Structural basis of membrane potential coupled vectorial CO_2_ hydration by the DAB2 complex in chemolithoautotrophs. Nature Communications 2026 17:1 17, 4071- (2026).

33. Kane, A. L. et al. Formate metabolism in *Shewanella oneidensis* generates proton motive force and prevents growth without an electron acceptor. J. Bacteriol. 198, 1337–1346 (2016).

34. Jormakka, M., Törnroth, S., Byrne, B. & Iwata, S. Molecular Basis of Proton Motive Force Generation: Structure of Formate Dehydrogenase-N. Science (1979). www.sciencemag.orgSCIENCEVOL2958 (2002).

35. Steinhilper, R., Höff, G., Heider, J. & Murphy, B. J. Structure of the membrane-bound formate hydrogenlyase complex from *Escherichia coli*. Nat. Commun. 13, (2022).

36. Lancaster, C. R. D. *Wolinella succinogenes* quinol:fumarate reductase—2.2-Å resolution crystal structure and the E-pathway hypothesis of coupled transmembrane proton and electron transfer. Biochimica et Biophysica Acta (BBA) - Biomembranes 1565, 215–231 (2002).

37. Pinske, C. & Sargent, F. Exploring the directionality of *Escherichia coli* formate hydrogenlyase: a membrane-bound enzyme capable of fixing carbon dioxide to organic acid. Microbiologyopen 5, 721–737 (2016).

38. Paul, S. et al. Architecture of the 8 MDa Hdr–Vhu–Fwd super-assembly in class I methanogens. Nature 2026 656:8129 656, 1054–1063 (2026).

39. Lampret, O. et al. Interplay between CN-Ligands and the Secondary Coordination Sphere of the H-Cluster in [FeFe]-Hydrogenases. J. Am. Chem. Soc. 139, 18222–18230 (2017).

40. Winkler, M. et al. A safety cap protects hydrogenase from oxygen attack. Nat. Commun. 12, (2021).

41. Stripp, S. T. et al. Second and Outer Coordination Sphere Effects in Nitrogenase, Hydrogenase, Formate Dehydrogenase, and CO Dehydrogenase. Chem. Rev. 10.1021/acs.chemrev.1c00914 (2022) doi:10.1021/acs.chemrev.1c00914.

42. Abramson, J. et al. Accurate structure prediction of biomolecular interactions with AlphaFold 3. Nature 2024 630:8016 630, 493–500 (2024).

43. The PyMOL Molecular Graphics System, Version 1.2r3pre, Schrödinger, LLC.

44. Jaenecke, J. et al. Subunit fusion unlocks rapid *in vitro* maturation for slowly activating heterodimeric [FeFe]-hydrogenases. Chem. Sci. 17, 7678–7689 (2026).

45. Akhtar, M. K. & Jones, P. R. Deletion of *iscR* stimulates recombinant clostridial Fe-Fe hydrogenase activity and H_2_-accumulation in *Escherichia coli* BL21(DE3). Appl. Microbiol. Biotechnol. 78, 853–862 (2008).

46. Li, H. & Rauchfuss, T. B. Iron carbonyl sulfides, formaldehyde, and amines condense to give the proposed azadithiolate cofactor of the Fe-only hydrogenases. J. Am. Chem. Soc. 124, 726–727 (2002).

47. Duan, J. et al. Crystallographic and spectroscopic assignment of the proton transfer pathway in [FeFe]-hydrogenases. Nat. Commun. 9, (2018).

48. Lampret, O. et al. Interplay between the CN-Ligands and the Secondary Coordination Sphere of the H-cluster in [FeFe]-Hydrogenases.

49. Stripp, S. T. In Situ Infrared Spectroscopy for the Analysis of Gas-processing Metalloenzymes. ACS Catal. 11, 7845–7862 (2021).

50. Van Dijk, C., Mayhew, S. G., Grande, H. J. & Veeger, C. Purification and Properties of Hydrogenase from *Megasphaera elsdenii*. Eur. J. Biochem. 102, 317–330 (1979).

51. Del Barrio, M. et al. Interaction of the H-Cluster of FeFe Hydrogenase with Halides. J. Am. Chem. Soc. 140, 5485–5492 (2018).

52. Senger, M. et al. Stepwise isotope editing of [FeFe]-hydrogenases exposes cofactor dynamics. Proc. Natl. Acad. Sci. U. S. A. 113, 8454–8459 (2016).

53. Winkler, M. et al. Accumulating the hydride state in the catalytic cycle of [FeFe]-hydrogenases. Nat. Commun. 8, (2017).

54. Albracht, S. P. J., Roseboom, W. & Hatchikian, E. C. The active site of the [FeFe]-hydrogenase from *Desulfovibrio desulfuricans*. I. Light sensitivity and magnetic hyperfine interactions as observed by electron paramagnetic resonance. Journal of Biological Inorganic Chemistry 11, 88–101 (2006).

55. Engelbrecht, V. et al. One isoform for one task? The second hydrogenase of *Chlamydomonas reinhardtii* prefers hydrogen uptake. Int. J. Hydrogen Energy 46, 7165–7175 (2021).

56. Odom, J. M. & Peck, H. D. Hydrogenase, electron-transfer proteins, and energy coupling in the sulfate-reducing bacteria *Desulfovibrio*. Annu. Rev. Microbiol. 38, 551–592 (1984).

57. Sahin, S. et al. Probing the ferredoxin:hydrogenase electron transfer complex by infrared difference spectroscopy †. 10.1039/d5sc00550g (2025) doi:10.1039/d5sc00550g.

58. Le Laz, S., Kpebe, A., Lorquin, J., Brugna, M. & Rousset, M. H_2_-dependent azoreduction by *Shewanella oneidensis* MR-1: Involvement of secreted flavins and both [Ni-Fe] and [Fe-Fe] hydrogenases. Appl. Microbiol. Biotechnol. 98, 2699–2707 (2014).

59. Ng, C. K., Cai Tan, T. K., Song, H. & Cao, B. Reductive formation of palladium nanoparticles by *Shewanella oneidensis*: Role of outer membrane cytochromes and hydrogenases. RSC Adv. 3, 22498–22503 (2013).

